# High prevalence of class 1 integrase and characterization of class 1 integron gene cassettes in multiresistant bacteria isolated from the gut microbiota of extended antibiotic treated *Salmo salar* fish farms

**DOI:** 10.1101/532663

**Authors:** Felipe Vásquez-Ponce, Sebastián Higuera-Llantén, Jimena Cortés, Natalia Zimin-Veselkoff, Sergio H. Marshall, Fernando O. Mardones, Jorge Olivares-Pacheco

## Abstract

The use of antimicrobials in aquaculture is a common practice. Chile is second larger producer of salmon worldwide, but unfortunately is the first consumer of antibiotics. Tonnes of florfenicol and oxytetracycline yearly are used in the Chilean salmoniculture to control the pathogens that threaten the sustainability of the industry. This excessive use of antibiotics have selected populations of resistant bacteria from the sediments and the water column that sorround the fish farms. In a recent work, our lab described the high prevalence of multiresistant bacteria and Antibiotic Resistance Genes (ARGs) in the gut microbiota of Antlactic salmon (Salmo salar) treated with high doses of antibiotics. In this work, we revisited the analysis of the previously described gut multiresistant bacteria grouped in banks of florfenicol resistant isolates (FB) and oxytetracycline resistant isolates (OB) looking for the presence of integron-integrase elements. These elements have been described as an important players in the Antimicrobial Resistance (AMR) phenomenon and they are considered a good markers of the anthopogenic activities pollution. The results showed that the 100% of the multiresistant isolates present the class 1 intagrase. Despite this result, no isolate from FB showed the typical structure of class 1 integrons: the presence in 3’-CS of *qacEΔ1/sul1* genes. While in OB, only 23% of the isolates showed this characteristic structure. Additionally, only four isolates of OB and none of FB showed recognisable gene cassettes and no genes of resistance to florfenicol and oxytetacycline appeared in them. Of these four isolates, three of them showed a single gene cassette containing the *dfrA-14* gene, which confers resistance to trimethoprim. Whilst the other isolate showed the *aac(6’)31*-*qacH*-*bla*_*oxa*2_ genes, which confers resistance to aminoglycosides, quaternary ammonium compounds and beta-lactams, respectively. Finally, it was possible to demonstrate that the described integrons probably come from anthropogenic activities like clinical settings and/or industrial animal husbandry, since they show integrases proteins identical to those carried by human pathogens.

## Introduction

According to the FAO report, Chile is the second largest producer of farmed salmon after Norway, but at the same time, it is the country that consumes the greatest amount of antimicrobials in this activity [1]. This is because the constant infectious diseases that lead to the death of millions of fish. The main disease affecting the three salmonid species farmed in Chile, Atlantic salmon (*Salmo salar*); Coho salmon (*Oncorhynchus kisutch*); and Rainbow trout (*Oncorhynchus mykiss*), is the Salmon Rickettsial Syndrome (SRS), caused by the facultative intracellular bacterium *Piscirickettsia salmonis* [2]. Although this pathogen has been described in all major salmon producing countries, such as Norway, Canada, Scotland, Ireland and Australia [3–5], both the genetic conditions of the Chilean isolates and the particular environmental conditions of the fjords of southern Chile make it much more aggressive than those described in other countries [2]. In addition to the extreme aggressiveness shown by this bacterium in Chile, none of the 40 vaccines available provide sufficient protection to prevent the development of the disease [6–8]. All of the above have led to the use of antibiotics as the main way to control this bacterium.

Between 2007 and 2017, almost 5500 tonnes of antibiotics have been used as an active substance, reaching an average of 500 g of antibiotic per tonne of salmon produced [9]. According to the latest report on the use of antimicrobials by the Servicio Nacional de Pesca y Acuicultura (*National Fisheries and Aquaculture Service*) (Spanish acronym: Sernapesca), 393 tonnes of antibiotics were used in 2017, of which 92.2% corresponded to florfenicol, 6.7% to oxytetracycline and 1% to flumequine [9]. The main route of administration of these antibiotics is through medicated food [10,11]. The administration of medicated food fundamentally affects the gut microbiota of the fish [12], since the constant exposure to antimicrobials leads to the selection of resistant bacteria and the increase of horizontal gene transfer (HGT) of those elements containing Antibiotic Resistance Genes (ARGs) [13]. In a recent article published by our laboratory, we have described that subsequent treatments with antibiotics select multiresistant bacteria with a high prevalence in ARGs both to florfenicol and oxytetracycline in the gut microbiota of fish [14].

One of the most important elements in the dispersal capacity of the ARGs are the Integron-integrases systems [15]. These elements are bacterial genetic platforms that allow the acquisition, storage, cleavage and rearrangement of genes located in mobilizable elements called gene cassettes [16]. The integron structure is formed by; i) an integrase, whose function is recombine circularized DNA known as gene cassettes; ii) a recombination site called *attI* and; iii) a promoter, PC, that control the genetic expression of the captured genes [17,18]. Integrons participate actively in the bacterial evolution and they are vehicles of gene exchange between the environmental resistome and commensal and pathogenic bacteria [19]. According to the amino acid sequence of integrase proteins, the integrons have been classified into 5 classes [15], however, only the class 1, 2 and 3 integrons are highly associated with the successful dispersion of the ARGs [20]. Even more the class 1 integrons are, precisely, the most described in pathogenic bacteria from humans and animals and, in turn, they are the most abundant integrases in the clinical environment since most of them show ARGs giving resistance to a wide variety of antimicrobials [19]. Thanks to these characteristics, class 1 integron-integrases elements have recently been proposed as indicators of pollution by Antibiotic Resistance Bacteria (ARB), ARGs, and other anthropogenic pollutants [21]

Taking into account the above mentioned facts, the main objective of this work is to determine the prevalence of class 1, 2 and 3 integron-intagrases in banks of bacteria resistant to florfenicol and oxytetracycline, which were obtained from the gut microbiota of fish coming from four fish farms subjected to treatments with high doses of antibiotics. Moreover, we have been able to demonstrate that these class 1 integron-integrases come from the other anthropogenic activities like clinical settings or lan industrial animal husbandry different to the aquaculture and that the constant exposure to antibiotics allows them to remain in the salmon farming system in Chile. The presence of these elements, indicators of contamination by human activities, in the gut microbiota of fish, make this system a perfect environment for the exchange of ARGs between environmental bacterial and fish commensal bacteria. Finally, these genetic elements could be easily released to the environment through the faeces of the fish.

## Materials and Methods

### Banks of resistant bacteria to florfenicol and oxytetracycline

For this study, characterized banks of resistant bacteria to florfenicol and oxytetracycline isolated in our lab were used [14]. Shortly, four Atlantic salmon (*Salmo salar*) fish farms were chosen, located in the Aysén Region, North Patagonia, Chile. The farms were selected because at the time of the sample all of them had applied more than one medicated food treatment with antibiotics. Bacteria were isolated from the faeces and the intestine of the fish and they were plated in TSA medium and incubated at 15, 25 and 37 °C. Minimal Inhibitory Concentration (MIC) to florfenicol and oxytetracycline for all isolates were estimated. Those isolated showing a MIC ≥128 μg/mL for florfenicol and ≥32 μg/mL for oxytetracycline were considered to be resistant bacteria, according to EUCAST clinical standards for enterobacteria group. Both banks were taxonomically characterized by amplification and sequencing of the 16S gene. The bank of bacteria resistant to florfenicol (FB) consists of 47 isolates, while the bank of bacteria resistant to oxytetracycline (OB) consists of 44 isolates [14]. This study was carried out in accordance with law 20,380 regarding animal welfare, as set out by the Chilean Health Ministry in the use of wild or protected animal species in biomedical research and approved by the National Fisheries Service (SERNAPESCA) and the Pontificia Universidad Católica de Valparaíso Bioethical Committee.

### DNA extraction and manipulation

Total DNA of all isolates was extracted using the AxyPrep™ Bacterial Genomic DNA Miniprep Kit (Axygen Biosciencies, USA) following the manufacturer’s instructions. All PCR products were purified from the gel using the UltraClean^®^ GelSpin^®^ DNA extraction kit (MoBio, USA). All PCR products were cloned using the pCR2.1 TOPO™ TA plasmid (Invitrogen, USA). All plasmids were extracted with the E.Z.N.A.™ Pasmid mini kit Omega Bio-Tek, USA).

### Detection of class 1, 2 and 3 integrases in resistant banks of bacteria

In order to determine and characterize the presence of integron-integrase elements in the banks of bacteria resistant to florfenicol and oxytetracycline, partial sequences of the genes which encode for integrase proteins *int1*, *int2* and *int3* were amplified. With this purpose, the primers Int1F-Int1R [22], Int2F-Int2R and Int3F-Int3R [23]were used. Primer used in this study are collected in table 1.

### Characterization of class 1 integrons

In order to characterize the complete structure of the class 1 integrons, the *sul1* and *qacEΔ1* (3’-CS) genes were amplified, using specific primers in the class 1 integrase positive isolates. Hence, the primers Sul1F-Sul1R [24] and qacEΔ1F-qacEΔ1R [25] were used. Furthermore, and in order to amplify gene cassettes contained in the variable region, the primers Hep58 and Hep59 were used according to White et al 2000 [26]. The sequence of the primers and the PCR conditions are collected in Table 1. All PCR products were cloned in the pCR2.1 TOPO™ TA (Invitrogen, USA) plasmid and sequenced. All sequences were analysed using the BLAST program [27] and the variable regions were assembled using the CLC Genomics Workbench 10 (QIAGEN). The complete sequence of the integrons were deposited in the GenBank database. Accession numbers of the complete class 1 integrons described are collected in table 4

**Table 1:**
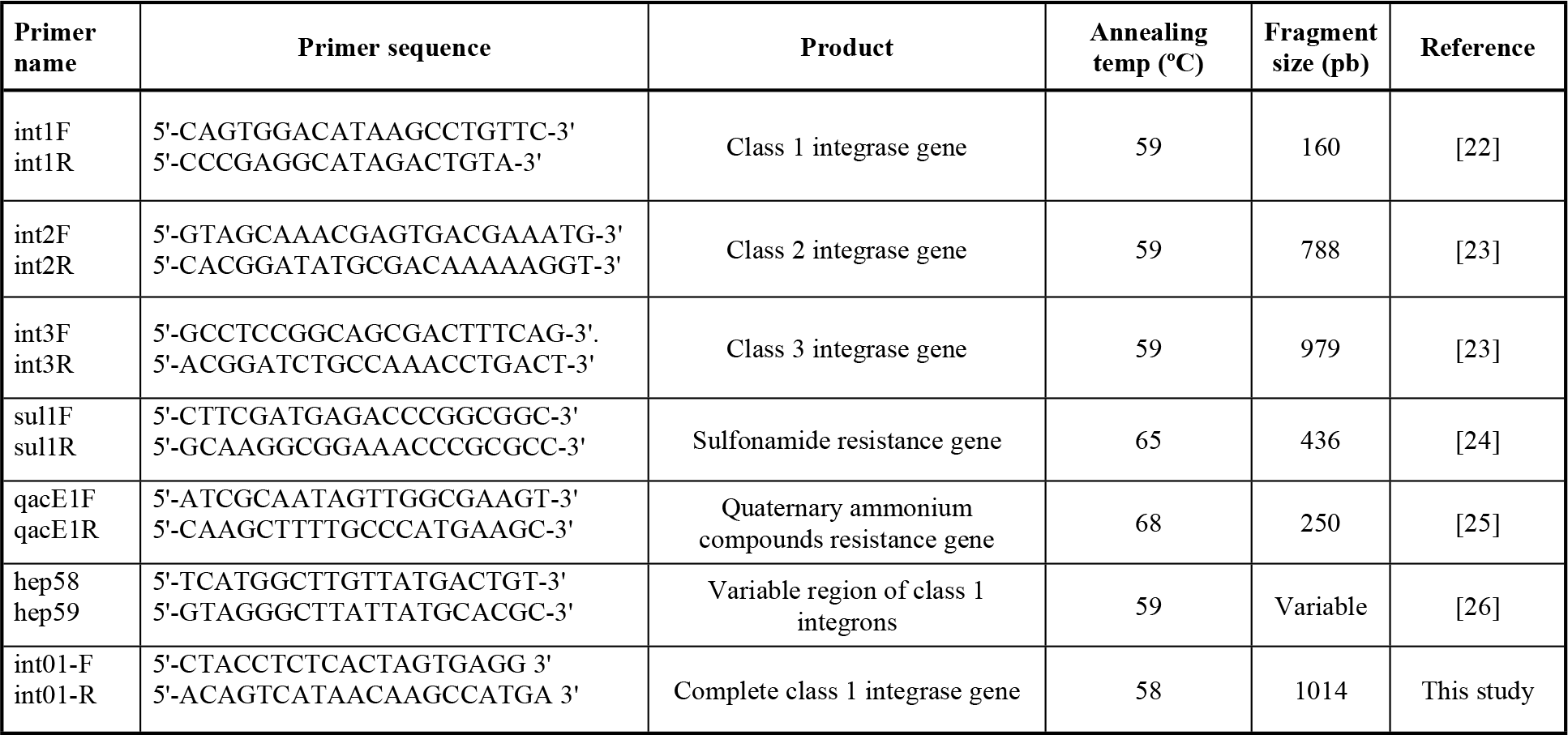
Primers used in this study

### Phylogenetic relationship of the class 1 integrase proteins

In order to identify the origin of class 1 integrase proteins, the full sequence was amplified using the primers int01-F and int01-R (Table 1). Furthermore, an extensive search for class 1 integrase proteins from different environments such as soil, fresh water, sea water and human pathogens were performed using the INTEGRALL database [28], with the purpose of identifying the phylogenetic origin of the integrase proteins from the banks of oxytetracycline and florfenicol. For the construction of the phylogenetic tree, the amino acid sequences from integrase proteins were used. The analysis of these sequences was performed using MEGA7 applying the maximum likelihood methods with 1,000 bootstraps. Accession number for the complete sequence of the integrases are collected in the supplementary table 1 (Table S1)

## Results

### Prevalence of integron-integrase elements in florfenicol and oxytetracicline isolate-banks

In order to determine the prevalence of the three types of integron-integrases elements, the search was carried out by PCR in both banks of resistant bacteria. We were able to identify that 100% of the isolates from both banks was positive for class 1 integrases (Tables 2 and 3). However, we were not able to find class 2 and 3 integrases. Furthermore, a search for the conserved elements of class 1 integrons of genes *sulI* and *qacEΔ*1 was made. One of the most important findings is that, despite all isolates of FB were positive for the class 1 intagrase, none of them was a carrier for the genes *sulI* and *qacEΔ*1 (Table 2). Likewise, 57% of the isolates showed the gene *sulI* in OB, while only 27% showed the gene *qacEΔ*1. Similarly, 23% showed both genes, which is the typical structure of a class 1 integron. Without a doubt, the most important feature of an integron is its variable region. Out of the 44 isolates of OB, only four were positive for the variable region (Table 3) and two isolates were found in different isolates of the same species *Serratia proteomaculans*. In the same way, another variable region was found in one of the important pathogens of salmon farming: *Aeromonas salmonicida*.

**Table 2:**
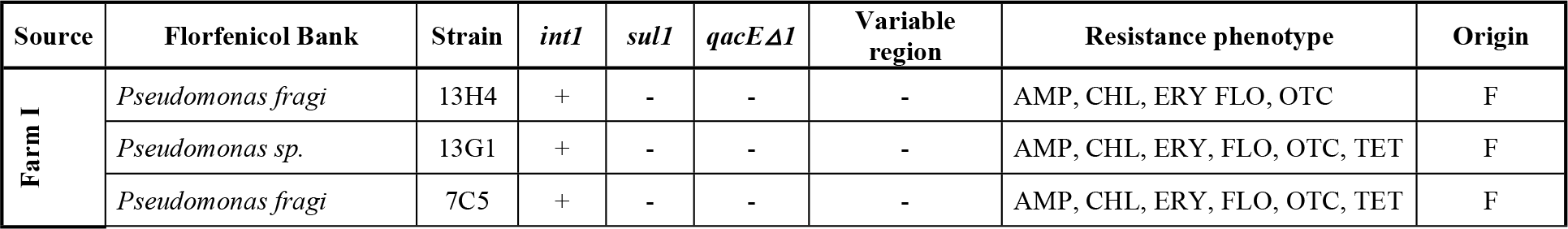
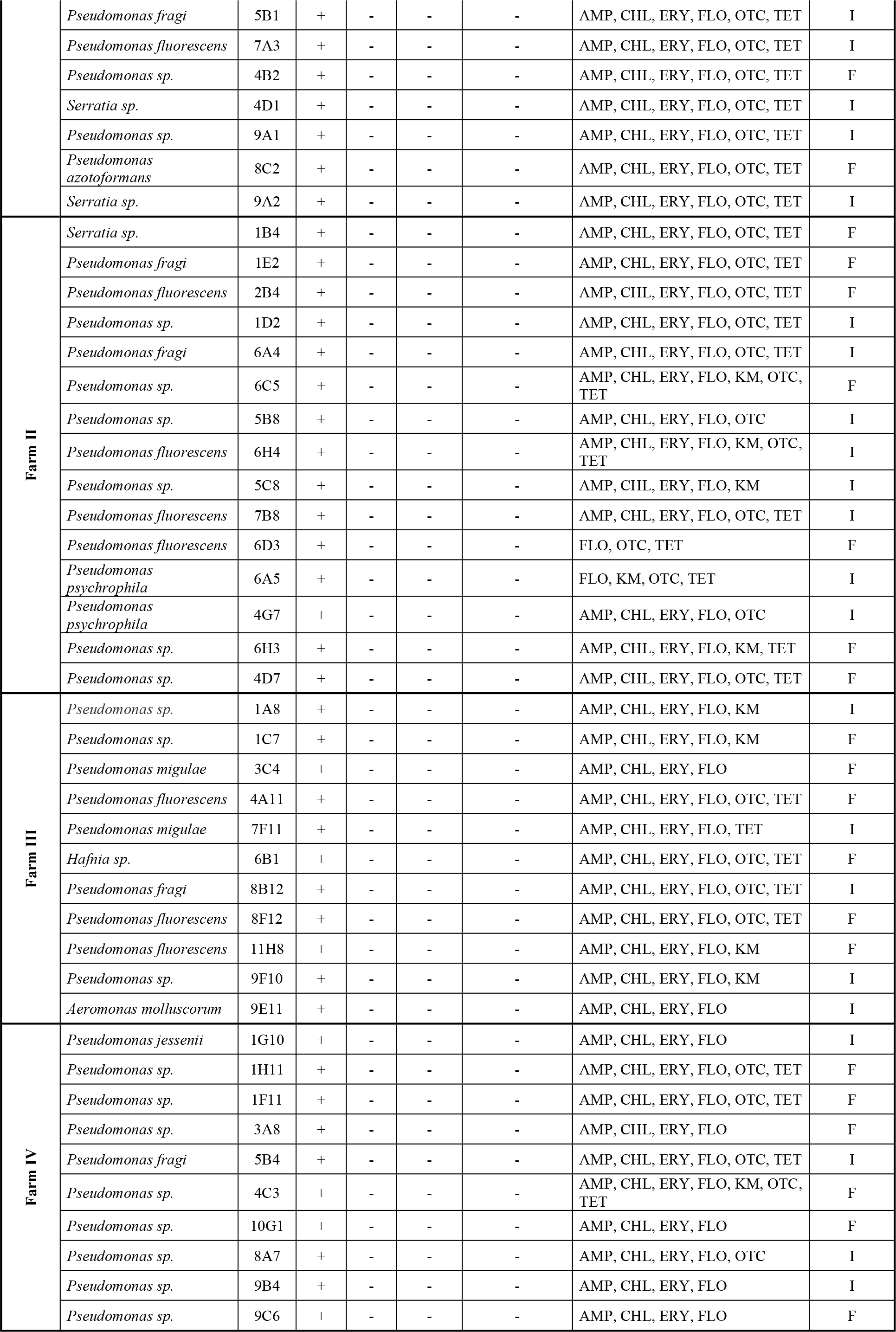
Florfenicol bank, antibiotic resistance profile and detection of class 1 integron elements. Antibiotics used, AMP (ampiciline), CHL (chloramphenicol), CIP (ciprofloxacin), ERY (erythromycin), FLO (florfenicol), KAN (kanamycin), TET (tetracycline). Resistance levels was defined by the EUCAST values for any antibiotic for enterobacteria group. AMP ≥64 μg/ml, CHL ≥32 μg/ml, CIP ≥4, ERY ≥16 μg/ml, FLO ≥128 μg/ml, TET ≥32 μg/ml. Origin: F= Fecal matter; I= Intestine.

**Table 3:**
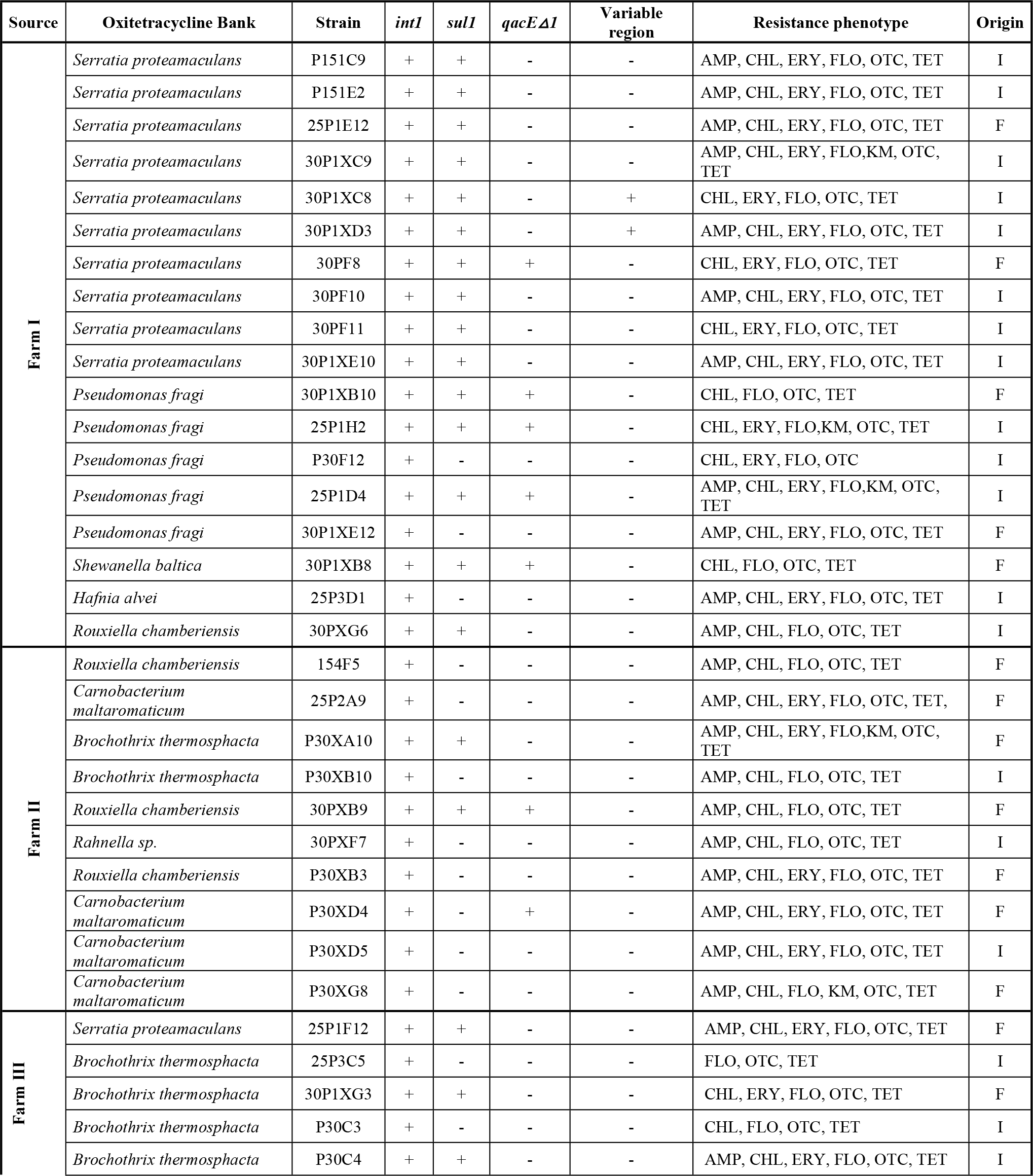
Oxytetracycline bank, antibiotic resistance profile and detection of class 1 integron elements. Antibiotics used, AMP (ampiciline), CHL (chloramphenicol), CIP (ciprofloxacin), ERY (erythromycin), FLO (florfenicol), KAN (kanamycin), TET (tetracycline). Resistance levels was defined by the EUCAST values for any antibiotic for enterobacteria group. AMP ≥64 μg/ml, CHL ≥32 μg/ml, CIP ≥4, ERY ≥16 μg/ml, FLO ≥128 μg/ml, TET ≥32 μg/ml. Origin: F= Fecal matter; I= Intestine.

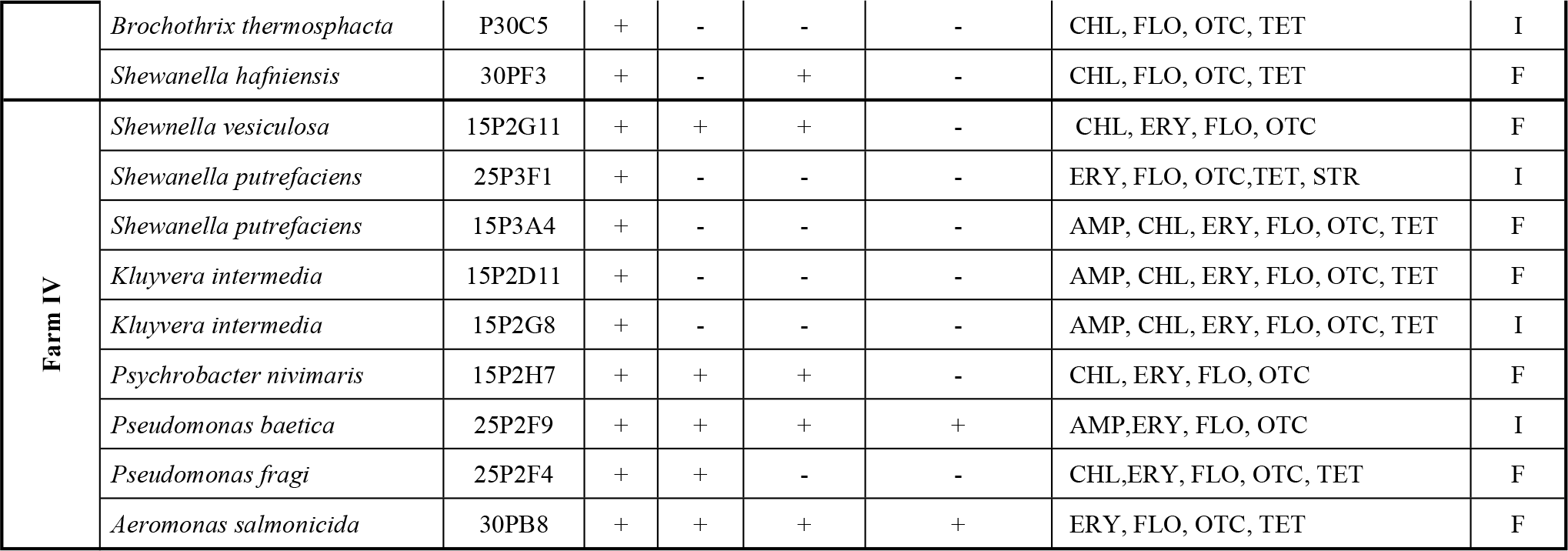

### Identification of gene cassettes in class 1 integrons

In order to identify possible resistance genes contained in the positive variable regions found in the bank of oxytetracycline resistant bacteria, the potential gene cassettes, amplified from the variable region, were sequenced and identified (Table 4). It was possible to identify gene cassettes in the strains *Pseudomonas baetica* 25P2F9 and *Aeromonas salmonicida* 30PB8, both isolated from fish farm IV. The gene cassette found in the isolate of *Pseudomonas baetica* contained the genes *aac(6’)31*-*qacH*-*bla*_*oxa-2*_ which encodes for both aminoglycoside adenylyltransferase - a protein resistant to quaternary ammoniums-, and for oxacillin-hydrolyzing class D beta-lactamase, respectively. With regard to *Aeromonas salmonicida* 30PB8, it was possible to identify the gene *dfrA14* which encodes for a trimethoprim resistance determinant, a dihydrofolate reductase enzyme. Finally, both isolates from the species *Serratia proteomaculans* showed the same gene cassette with the gene *dfrA14* (Figure 1). The other strains containing the genes *sulI* and *qacΔE*, in the typical structure of a class 1 integron, showed no gene cassettes in the variable region, which implies that the system has a great potential to incorporate and express new antibiotic resistance genes. Nonetheless, it is important to note that none of the antibiotic resistance genes to florfenicol and oxytetracycline described in both banks by Higuera-Llanten et al 2018 [14] are found in the integrons of the isolated bacteria. Currently, a complete metagenomic analysis is being carried out to find the relationship between these resistance determinants and these genetic elements that could probably be established in non-culturable bacteria.

**Table 3:**
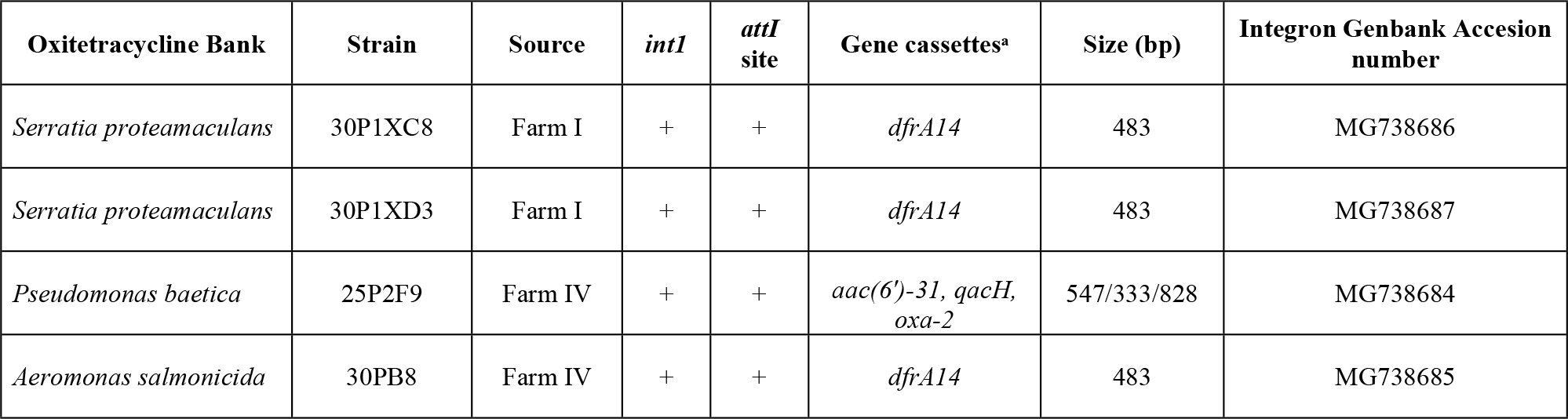
Gene cassette composition in positive Class 1 integrase variable region strains. Genes found: i) *dfrA14* (Trimethoprim-resistant dihydrofolate reductase), resistance to trimethoprim; ii) *aac(6’)-31* (Aminoglycoside adenylyltransferase), resistance to aminoglycosides; iii) *qacH* (Quaternary ammonium protein), resistance to quaternary ammonium compounds and; iv) *oxa-2* (Oxacillin-hydrolyzing class D beta-lactamase), resistance to beta-lactam antibiotics.

**Figure 1.**
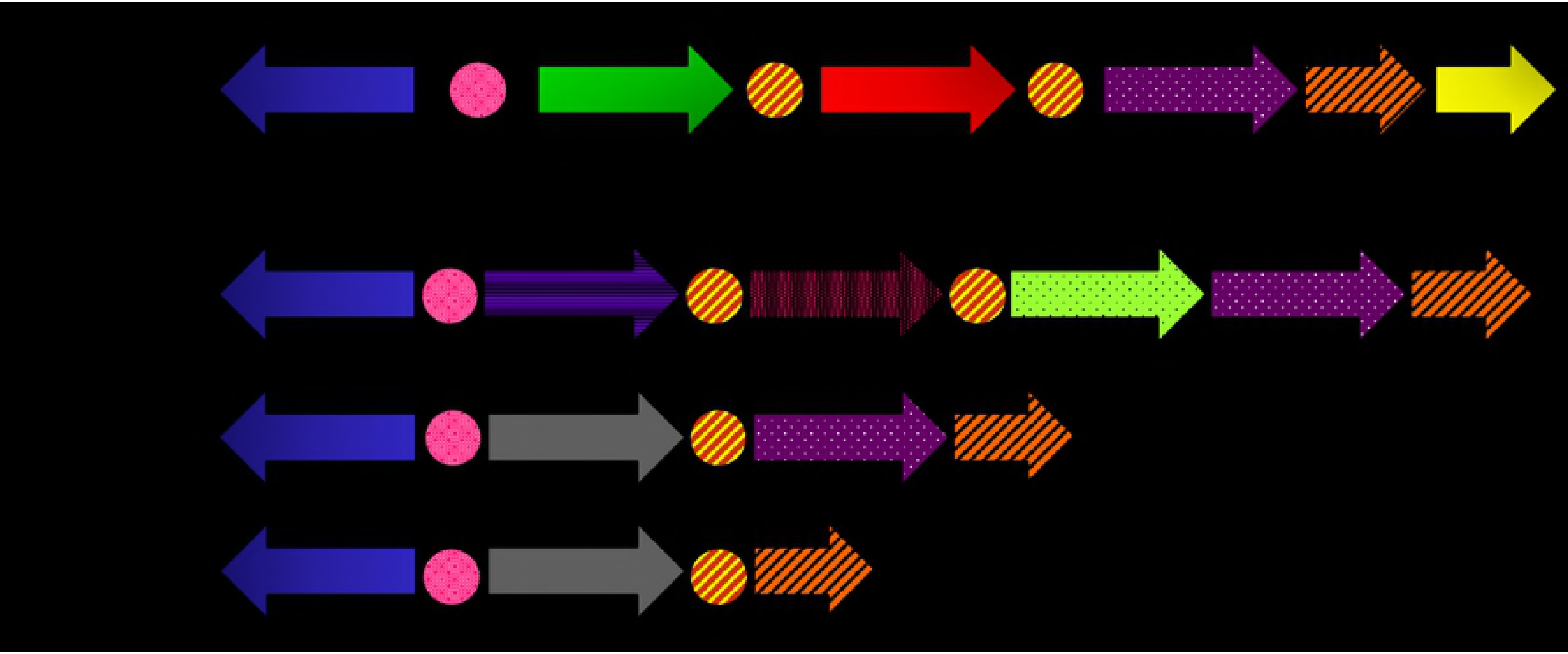
Scheme of Class 1 integrons. A) Basic class 1 integrons platform. 5’ CS: 5’- conserved segment consists of *intI* (gene which encodes for integrase), Pc (promotor), *attI* (recombination site). Structured gene cassettes formed by the recombination site *attC*. 3’ CS: 3’-conserved segment consists of the genes: i) *qacEΔ1* (resistance to quaternary ammonium); ii) *sul1* (resistance to sulfonamides) and; iii) *orf5*, whose function is completely unknown. **B) Integrones found in the microbiota of *Salmo salar***.

### Class 1 integrases isolated from gut microbiota of *Salmo salar* show a close phylogenetic relationship with clinical important bacteria

In order to evaluate the phylogenetic relationship and, therefore, the possible origin of class 1 integrases identified in the gut microbiota of *Salmo salar*, the complete ORF of 23 genes of the class 1 integrase from both banks was obtained, which were translated into their amino acid sequence. It is worth noting that it was not possible to obtain the gene *intI1* in all the isolates with the primers used. Despite this, a phylogenetic analysis was carried out with the 23 integrases of the isolates from both banks of resistant bacteria, along with various integrases obtained from different environments such as sea water, soil, fresh water and human pathogens. In the obtained phylogenetic tree (Figure 2), 3 differential clades are clearly shown, where integrases from marine environment, soil, fresh water, and from human pathogens are identified. The 23 integrases from the isolates of gut microbiota of *Salmo salar* are grouped in the clade corresponding to human pathogens of clinical importance, such as *Acinetobacter baumannii*, *Salmonella enterica*, *Enterobacter cloacae*, *Pseudomonas aeruginosa* and *Klebsiella pneumoniae*. These data suggest that there is an important contribution component from land anthropogenic activities like clinical settings and/or animal husbandry to the gut microbiota of salmon, and the antibiotics used in this productive activity select indirectly these elements keeping them and dispersing them in the marine environment.

**Figure 2.**
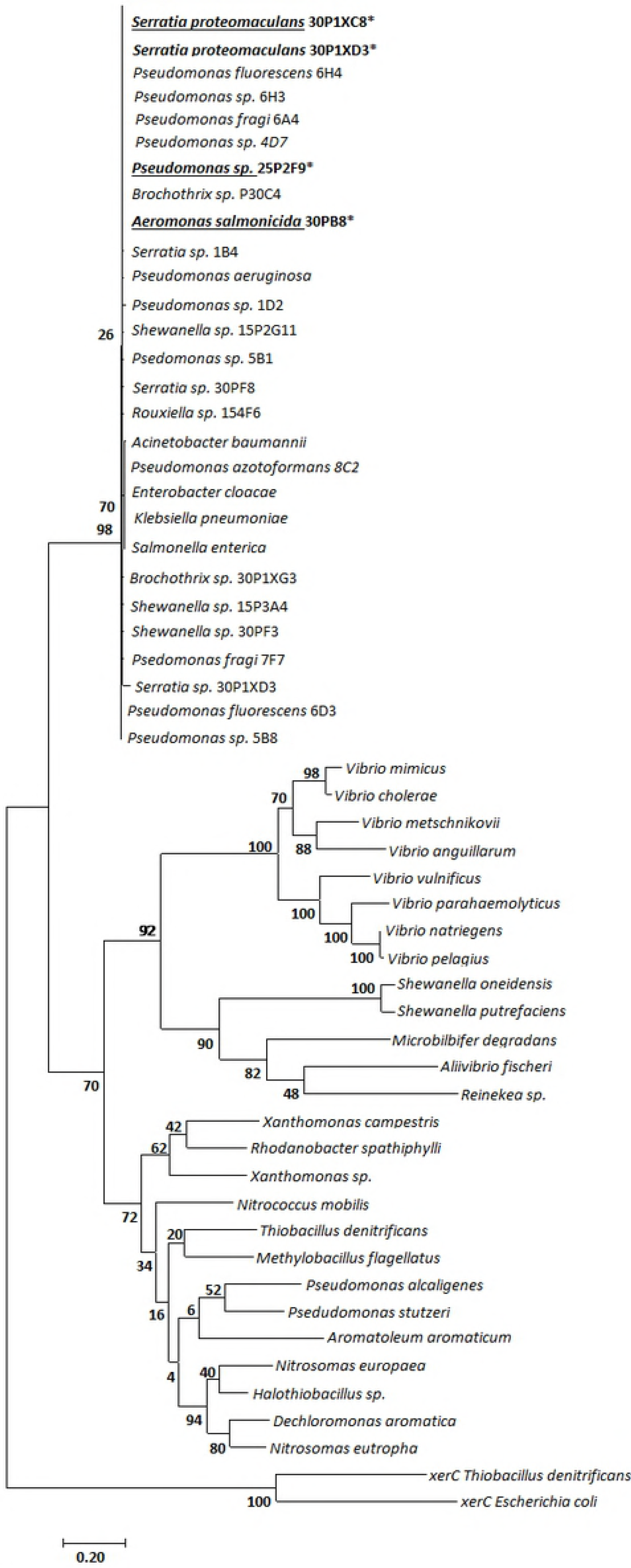
Phylogenetic relationship of class 1 integrases from the salmon industry in Chile. The phylogenetic tree shows class 1 integrases from diverse environments such as seawater, soil, fresh water and human pathogens. It is clearly observed that the integrases from the salmon farming in Chile are grouped together with integrases of pathogens of clinical importance and marked with the Δ symbol. XerC proteins from *Escherichia coli* and *Thiobacillus denitrificans* were used as outgroup. The results were obtained using the maximum likelihood methods after 1,000 bootstraps.

## Discussion

In this work, the high prevalence of class 1 integron-integrase elements in banks of bacteria resistant to florfenicol and oxytetracycline, isolated from the gut microbiota of the Atlantic salmon (*Salmo salar*) treated with high doses of antibiotics in salmon farms has been reported. In a previous laboratory work, it was demonstrated that the high use of antimicrobials brings with it a high abundance of multiresistant bacteria with a high presence of ARGs to both florfenicol and oxitetrecycline. More than 50% of the isolates, counting both banks, showed resistance against at least 4 of the seven antibiotics tested. Despite the fact that an extensive work was performed, in this last work, the presence of integron-like elements was not measured. These genetic elements have been considered of great importance in the antimicrobial resistance phenomenon, since they could play a fundamental role in the transfer of genes between the clinical and the natural environment [16,19]; hence, it has even been estimated that its role in the marine environment is much more important than in the terrestrial environment [29].

The occurrence of the three classes of integrons-integrases in the environment has been studied in different works, especially the abundance of class 1 integrons-integrases and the composition of their respective gene cassettes. In this study, the presence of the three classes of integrons-integrases so far described Classes 1, 2 and 3 was measured, and it was found that 100% of the isolates from both banks showed class 1 integrons; however, no isolate showed class 2 and class 3 integrons. This type of result had already been described in salmonid production systems. In salmon farms of rainbow trout in Australia, 31% of the isolates of *Aeromonas* spp. showed class 1 integrons. Nevertheless, class 2 and class 3 integrons were not detected [30]. The same result was obtained with isolates from *Pseudomonas* spp., where 23% was positive for class 1 integrons, while class 2 and class 3 integrons were not detected either [31]. The same was found in catfish farming, where 33% of the isolates from *Pseudomonas* and 28% of the isolates from *Aeromonas* showed class 1 integrases, but all of them were negative for class 2 and class 3 intregrases [32]. Only in confined eel farming, in China, the presence of the three classes of integrons in systems related to aquaculture has been demonstrated [33]. Although class 2 integrons-integrases have been detected frequently in different types of environments, class 3 integrons-integrases have rarely been detected outside the clinical setting [16,33]. The high frequency of the appearance of class 1 integrons lacking the typical structure of the 3’-CS is another interesting result of this work, since only 23% of OC showed the genes *qacE*Δ1/*sulI*, while none of the isolates of FB, showed these genes in its structure.

But without a doubt, the most important result of this work is that despite the fact that all bacteria showed a high prevalence of antibiotic resistance genes to florfenicol and oxytetracycline, none of these genes were part of these genetic elements. Although tetracycline resistance genes are found with a high frequency in all environments impacted by human activity [34], their presence in class 1 integrons is very scarce. Only sometimes the genes *tetC* and *tetE* have been found to be associated with class 1 integrons [35]. However, due to the high amount of antibiotics used in this industry and the high volume of resistance genes present in the obtained resistant isolates, there is a tendency to think that the incorporation of these genes in this type of gene elements could be favoured, but it seems that the dynamics of the integron-integrases elements in this system is not so evident. A similar case occurs with the genes resistant to florfenicol, *floR* and *fexA.* Up to now, the gene *fexA* has never been described in integrons, while the gene *floR* has rarely been found in class 1 integrons. This is mainly because they are more directly related to horizontal transfer elements such as plasmids and transposons [33]. This fully confirms that despite a high incidence of the genes *floR* and *fexA* in the resistant bacteria isolated from the fish’s microbiota, none of these genes could be found as a part of the integrons.

Although the 91 resistant isolates present in both banks were positive for the presence of class 1 integrase, only four of them showed recognisable gene cassette structures. In all isolates, it was possible to find genes that appear with high frequency in class 1 integrons. Such is the case of the cassette array found in the species *Pseudomonas baetica*, where the genes *aac(6’)-31*, *qacH* and *bla*_*oxa-2*_ are found. The first encodes for aminoglycoside adenylyltransferase; these genes appear with a very high frequency in class 1 integrons [16] since it has been described that more than 80% of these elements contain these types of enzymes in their structures [36]. Thus, it has even been described that genes resistant to aminoglycosides are permanent elements in integrons described in the marine environment [29,37] which gives even more dynamism to the antibiotic resistance phenomenon in aquatic environments [38]. Regarding the gene *qacH*, which encodes for a quaternary ammonium resistance mechanism, it was first described in members of the genus *Staphylococcus* [39] but, nowadays, it is frequently found in Gram-negative bacteria, giving them resistance to a broad spectrum of quaternary ammonium compounds, unlike for other genes *qac* [40,41]. Another characteristic of this gene is that it appears relatively often in the food contaminating species *Listeria monocytogenes*, which is associated with resistance to disinfectants derived from benzalkonium chloride [42,43]. This element of resistance has also been found in other bacteria that are frequent food contaminants, such as the members of the genus *Salmonella* [44] and *Staphylococcus aureus*, which contaminates milk [45]. Considering that salmon meat is intended for human consumption, the presence of this element of resistance could represent a risk for eliminating this type of bacteria from the processing plant.

Unquestionably, beta-lactamases are the most studied elements in the antibiotics resistance phenomenon, since they represent the biggest problem for human health. OXA-type beta-lactamases have been widely characterised and more than 350 different alleles have been described worldwide [46]. OXA-type beta-lactamases confer resistance to any type of penicillin and some of them can extend their spectrum of activity to cephalosporins [47]. Most of them are part of plasmids in Gram-negative bacteria and they appear relatively often in class 1 integrons, exclusively in human pathogens such as *Pseudomonas aeruginosa* [48], *Salmonella tiphymurium* [49], *Escherichia coli* [50], and *Acinetobacter baumanii* [51]. Those, that have been detected outside the clinical environment, have always been detected in pathogenic bacteria or human commensals [52,53]. Therefore, this is the first time that an OXA-type beta-lactamases betalactamasa, taking part of a class 1 integron associated with aquaculture bacteria, is described.

Regarding the integrons found in *Aeromonas salmonicida* and *Serratia proteomaculans*, in both cases, the resistance gene to trimethoprim *dfrA14* has been found, which encodes for one of the most distributed alleles of a dihydrofolate reductase. The association of integrons with genes which encode for dihydrofolate-reductases has been widely documented in aquaculture [30,33,54], it has even been possible to demonstrate the presence of these elements in marine sediments associated with the production of salmon in Chile [55,56].

Integrons are widely distributed elements in all environments; however, based on the in the IntI Integrase protein sequence, a classification that has allowed them to be classified into different types has been established, and it has been demonstrated that some of them are much more frequent depending on the environment in which they are found [16]. This is how integrases typical of soil bacteria and fresh water, marine bacteria and pathogenic bacteria, both veterinary and human, have been classified [17]. It was expected that the bacteria present in the fish’s microbiota would be rich in integrons, either from freshwater, from its initial breeding, or from seawater bacteria. Nevertheless, all sequenced integrases are those normally found in human pathogens. This suggests that these elements have come from the land anthropogenic activities like clinical settings and/or animal husbandry and have been highly successful in the salmon farming environment, which could be dangerous due to the probability of exchange of ARGs between land and marine systems.

The high presence of integron-integrases elements in aquaculture has been described in several papers [30,33,57,58], even further, the presence of integrons in resistant bacteria isolated from the sediment of salmon farms in Chile has been described in a recently published paper by [56] which has even reached clinical isolates of the bacteria *E. coli*. Among the gene cassettes described in this work, the high abundance of genes *dfrA12* and *dfrA17*, which confer resistance to trimethoprim, as well as genes resistant to aminoglycosides *aadA2* and *aadA5*, can be observed. These genes appear with high frequency in the integron-integrases elements; however, none of them contained genes for resistance to florfenicol or oxytetracycline either, which are the most used antimicrobials in salmon farming in Chile. These results are fully validated in this work.

With the data obtained in the present study, we can conclude that the presence of integron-integrase elements is highly abundant in the gut microbiota in farmed Atlantic salmon subjected to treatments with high doses of antimicrobials florfenicol and oxytetracycline. In addition to this, we can say that these elements apparently come from land human activities like clinical settings and/or animal husbandry, since the class 1 integrase gene is identical to that found in human pathogenic bacteria of clinical importance such as *P. aeruginosa* or *A. baumanii*. Thus, the contribution of the human activity is could be the main cause of dispersion and dissemination of ARGs in natural environments; also, the large amount of antimicrobials used in aquaculture only favours the maintenance and perpetuation of these elements in the environment. This becomes even more delicate since, if we consider the gut microbiota of fish as a "semi-isolated" system from the environment and due to its direct contact ability with the used antibiotics, it could become the perfect place for genetic exchange to occur between bacteria from different environments. Hence, the gut microbiota of fish treated with high doses of antibiotics could become an ideal reservoir for ARGs, which have a high probability of being dispersed through the faeces, loading the marine environment with these types of genetic elements. The high use of antimicrobials requires a quick solution, consequently, the different production companies are already beginning to take measures to reduce the use of antimicrobials, in the hope that by the year 2020, the use of these drugs is reduced by at least 50%.

## Acknowledgements

This study was supported by CONICYT-Chile through the project FONDECYT 11150858 and by the interdisciplinary project Envirotracker 039.461/2017.

## Author Contributions

**Conceptualization:** Jorge Olivares, Felipe Vasquez

**Data curation:** Sebastián Higuera, Felipe Vásquez, Jimena Cortés

**Formal analysis:** Jorge Olivares, Sebastián Higuera, Felipe Vásquez, Fernando Mardones, Sergio Marshall, Natalia Zimin, Jimena Cortés

**Funding aquisition:** Jorge Olivares

**Methology:** Sebastián Higuera, Felipe Vásquez, Natalia Zimin

**Writing-Original draft:** Felipe Vasquez, Jorge Olivares, Jimena Cortés

**Writing-review & editing:** Jorge Olivares, Fernando Mardones, Sergio Marshall

